# Genome-wide survey and expression analysis of the *SLAC/SLAH* gene family in pear (*Pyrus bretschneideri*) and other members of the *Rosaceae*

**DOI:** 10.1101/288308

**Authors:** Guodong Chen, Xiaolong Li, Xin qiao, Jiaming Li, Li Wang, Xiaobing Kou, Xiao Wu, Guoming Wang, Hao Yin, Peng Wang, Shaoling Zhang, Juyou Wu

## Abstract

S-type anion channels (*SLAC/SLAH*s), which play important roles in plant anion (such as nitrate and chloride) transport, growth and development, abiotic stress responses and hormone signaling. However, there is far less information about this family in *Rosaceae* species. We performed a genome-wide analysis and identified *SLAC/SLAH* gene family members in pear (*Pyrus bretschneideri*) and four other species of *Rosaceae (Malus domestica*, *Prunus persica*, *Fragaria vesca* and *Prunus mume*). A total of 21 *SLAC/SLAH* genes were identified from the five *Rosaceae* species. Based on the structural characteristics and a phylogenetic analysis of these genes, the *SLAC/SLAH* gene family could be classified into three main groups (I, II and III). The evolutionary analysis showed that the *SLAC/SLAH* gene family was comparatively conserved during the evolution of *Rosaceae* species. Transcriptome data demonstrated that *PbrSLAC*/*SLAH* genes were detected in all parts of the pear. However, *PbrSLAC1* showed a higher expression level in leaf, while *PbrSLAH2/3* was mainly expressed in roots. In addition, *PbrSLAC/SLAH* genes were only located on the plasma membrane in transient expression experiments in *Arabidopsis* protoplasts cells. These results provide valuable information that increases our understanding of the evolution, expression and functions of the *SLAC/SLAH* gene family in higher plants.

## 1. Introduction

As we know, anions, such as nitrate, chloride, sulfate and phosphate, as well as organic anions, such as malate and citrate, are essential for the healthy growth and development of animals and plants [1]. Anions can not only be used as nutrients for animals and plants through absorption, such as nitrate ions being the main nitrogen source for plant growth, but also control the growth and development of animals and plants as signal molecules. Anion channels, aqueous pores formed by integral membrane proteins, regulate the transport of anions across the membrane [2]. In animals, studies have focused on the chloride anion channel and cystic fibrosis transmembrane conductance regulator (*CFTR*) anion channel. Recently, nine *CLC*-type chloride channels have been found that are mainly involved with the excitability of cells in vivo, transcellular transport of substances within the cells, and the regulation of pH values [3]. On the contrary, a variety of anion channels exist in plants and are divided into rapid anion channels (R-type) and slow anion channels (S-type) [4]. Among them, the R-type contains the *CLC* and *AMLT1* families, which act in the millisecond range and gate the membrane potential, having V-shaped current–voltage relationships [5]. However, the S-type only contains the *SLAC1* family, which have slow activation/deactivation kinetics in the 10 s range and weak voltage dependency values [6, 7]. Genome-wide analyses and comparative genomics provide insights into the structures, evolution and functions of gene family members [8]. The S-type anion channel contains five *SLAC/SLAH* gene family members, SLAC1, SLAH1, SLAH2, SLAH3 and SLAH4 proteins, which form the founder anion channel of a small gene family in the model plant *Arabidopsis* [9]. Recently, gene functional analyses of the *SLAC/SLAH* family were reported in rice (*Oryza sativa*) [10], poplar (*Populus trichocarpa*) [11], barley [12] and tobacco [13]. Interestingly, more investigations have focused on the functions of slow-anion channel genes, especially those involved in nitrate ion absorption and transport.

In *Arabidopsis*, previous studies have suggested that the *SLAC/SLAH* genes play important roles in stress signaling, growth and development, and hormone responses [14]. For instance, the *AtSLAC1* and *AtSLAH3* genes interactions with distinct kinase phosphatase pairs are related to water-stress signaling [15–18]. The *OsSLAC1* gene is essential for stomatal closure in the face of CO_2_-stress signaling [10, 19]. In addition, *SLAC1* is mainly distributed in guard cells, and it is activated by phosphorylation from the *OST1* kinase, leading to induced rapid stomatal closure [20]. For hormone signaling, *SLAC1* was activated by abscisic acid (ABA) through direct interactions with *OST1* and CPKs, such as *CPK3/6/21/23,* which helps to enhance drought tolerance by regulating stomatal closure [16]. *SLAC/SLAH*s are also involved in plant growth and development. For instance, there is evidence for the Ca^2+^-dependent *CPK2/CPK20* regulation of the anion channel *SLAH3* gene that regulates pollen tube growth in *Arabidopsis* [21]. Moreover, many studies of the *SLAC/SLAH* gene family have also been carried out in other species, such as barley [12], poplar [11] and tobacco BY-2 cells [13].

The functions and characteristics of the *SLAC/SLAH* gene family have been investigated in the model plant *Arabidopsis*. For example, *SLAC1* is mainly expressed in guard cells and weakly in root/stamens/young siliques. *SLAH3* is found in mesophyll cells and shows weaker expression levels in guard cells and stamens, but is strongly expressed in roots. *SLAH2* is the closest homolog of *SLAH3*, and *SLAH1* is expressed in both the root and hypocotyl/stamens. *SLAH4*, which is phylogenetically most closely related to *SLAH1*, shows a stronger expression near the root tip [9, 18]. Considering the expression patterns of *SLAC/SLAH* family members in roots and the nitrate-permeable channel activity reported earlier [18], we hypothesized that some members of the *SLAC/SLAH* gene family may be involved in nitrate fluxes and translocation in *Rosaceae*. For instance, *SLAC1* and *SLAH3* display nitrate transport activities when expressed in oocytes with NO_3_^−^/Cl^−^ permeability ratios of 10 and 20, respectively [18]. Compared with *SLAC1*, *SLAH3* exhibits a higher preference for nitrate [16], and *SLAH3*, a nitrate efflux channel, plays a key role in the nitrate-dependent alleviation of ammonium toxicity in plants [9]. *SLAH2* is expressed in the immediate surroundings of the vasculature, and these cells determine the anion composition of the sap flow between roots and shoot [22]. These results provide strong supports for the analysis of these genes in the *Rosaceae* species. In this study, we employed bioinformatics and publicly available data to identify the *Rosaceae SLAC/SLAH* genes on a genome-wide scale and mainly analyzed their functions in pear. Our study aimed to annotate the full-length *SLAC*/*SLAH* genes in pear and other *Rosaceae* species, as well as to explore their subcellular localization and examine their expression levels in different pear tissues. We investigated shortages or excesses in related nutrients in response to nitrate, and provide a relatively complete profile of the *SLAC/SLAH* gene family in *Rosaceae*. More significantly, a comparison of *SLAC/SLAH* gene structures, evolution, and experimental data between *Arabidopsis* and the *Rosaceae*, provides insights into the functions of the *Rosaceae SLAC/SLAH* genes. Our results provide a set of potential candidate *SLAC/SLAH* genes for the future genetic modification of anion transport and stress tolerance in higher plants.

## 2. Materials and Methods

### 2.1 Identification of *SLAC/SLAH* gene family members in pear and other Rosaceae species

To identify the *SLAC/SLAH* genes in pear and other *Rosaceae* species, multiple database searches were performed. The *Arabidopsis* SLAC/SLAH protein sequences AtSLAC1 (AT1G12480), AtSLAH1 (AT1G62280), AtSLAH2 (AT4G27970), AtSLAH3 (AT5G24030) and AtSLAH4 (AT1G62262) were downloaded from the *Arabidopsis* Information Resource (TAIR) (http://www.arabidopsis.org/) [23] and the rice SLAC/SLAH protein sequences Os04g48530.1, Os01g43460.1, Os05g13320.1, Os01g28840.1, Os01g12680.1, Os07g08350.1, Os05g50770.2, Os01g14520.1 and Os05g18670.1 were downloaded from Phytozome (http://phytozome.jgi.doe.gov/pz/portal.html#). These sequences were used as query to perform BLAST algorithm-based searches against pear and other *Rosaceae* genome databases. The strategy to acquire every gene of the *SLAC/SLAH* family was the following: The pear (*Pyrus bretschneideri*) genome sequence was downloaded from the pear genome project (http://peargenome.njau.edu.cn/) [24]. The genome sequences of apple, peach and strawberry were downloaded from Phytozome. The Chinese plum genome sequence was downloaded from the *Prunus mume* Genome Project (http://prunusmumegenome.bjfu.edu.cn/index.jsp). Additionally, the seed alignment file for the *SLAC/SLAH* domain (PF03595.13) obtained from the Pfam database was used to build a Hidden Markov Model (HMM) file using the HMMER3 software package [25, 26]. HMM searches were then performed against the local protein databases of pear and other *Rosaceae* species using HMMER3. Furthermore, all obtained *SLAC/SLAH* protein sequences were analyzed against the Pfam database to verify the presence of *SLAC1* domains. The *SLAC1* domain was also detected by the SMART program (SMART:http://smart.embl-heidelberg.de/). Protein sequences lacking the *SLAC1* domain or having E-values more than 1e-6 were removed.

### 2.2 Structure of the *SLAC/SLAH* genes and a conserved motif analysis

The structures of the *SLAC/SLAH* genes were analyzed using Gene Structure Display Server (GSDS 2.0) (http://gsds.cbi.pku.edu.cn/) by aligning the cDNA sequences with their corresponding genomic DNA sequences. Conserved motifs of the SLAC/SLAH proteins were identified using the online Multiple Expectation Maximization for Motif Elicitation (MEME) (http://meme.nbcr.net/meme/cgibin/meme.cgi) [27].

### 2.3 Phylogenetic analysis

The phylogenetic trees were constructed based on the SLAC/SLAH protein sequences from pear, rice, *Arabidopsis* and the other members of *Rosaceae* using the Neighbor-Joining (NJ) method in MEGA6.0 (http://www.megasoftware.net/) [28], and the bootstrap test was carried out with 1,000 replicates. The p-distance and the pairwise deletion option parameters were selected.

### 2.4 Chromosomal localization and synteny analysis

The chromosomal localization information of the *Rosaceae SLAC/SLAH* was obtained from genome annotation files (http://peargenome.njau.edu.cn). Then, a method similar to that developed for the PGDD (http://chibba.agtec.uga.edu/duplication/) [29] was used to perform the synteny analysis. Initially, local all-vs-all BLASTP algorithm-based searches between pear and the other four *Rosaceae* species, and between pear and *Arabidopsis* genomes, were conducted to identify potential homologous gene pairs (E <1 e^−10^). Afterwards, MCScanX was used to identify syntenic chains using homologous pairs as input [30]. The downstream analysis tools in the MCScanX package were used to identify whole-genome (WGD)/segmental, tandem, proximal and dispersed duplications in the *SLAC/SLAH* gene family. The results were displayed using Circos software [31].

### 2.5 Subcellular localization of the *SLAC/SLAH* genes of pear and *Arabidopsis*

The full-length coding sequences of the *SLAC/SLAH* genes of pear and *Arabidopsis*, including *PbrSLAC1*, *PbrSLAH2/3-1*, *PbrSLAH2/3-2* and *PbrSLAH2/3-3* and *AtSLAC1*, *AtSLAH1*, *AtSLAH2*, *AtSLAH3* and *AtSLAH4* were amplified from pear and *Arabidopsis* roots, respectively. The amplified PCR products were digested with XbaI and BamHI, and cloned directionally into the modified PC1301 vector containing the CaMV 35S promoter (Clontech, Beijing, China). These manipulations resulted in the p*SLAC/SLAH*s-GFP. Primers used for cloning genes and for constructing vectors are shown in Supplementary table S3. Mesophyll protoplasts from *Arabidopsis* were isolated [32] and transformed [33] as previously described [34]. Images were processed using the Zeiss LSM Image Browser (Zeiss LSM 780, Germany). Finally, at least three independent transient expression assays were performed for each construct.

### 2.6 Quantitative real-time PCR

Roots were harvested from five-week-old pear seedlings grown under controlled growth conditions. They were then grown in hydroponic nutrient solutions containing different concentrations of nitrate ions for two weeks. Total RNA was isolated from the collected pear tissues using an RNA kit (RNA simply Total RNA Kit, Tiangen, and Beijing, China) according to the manufacturer’s instructions. Then 3 μg of each sample was reverse transcribed into cDNA using the PrimeScript RT reagent Kit (Trans Gen). The specific primers of the *SLAC/SLAH* genes of pear and *Arabidopsis*, and the housekeeping actin gene (Pbr035825.1), were designed using the Primer Premier 5.0 software (Supplementary Table S3). To verify the specificity of these primers, we used the program Primer search-Paired against the pear genome. The qRT-PCR assays were performed with three biological and three technical replicates. All of the real-time PCR reactions included 2 μl diluted cDNA, 200 nM of each primer, 2× SYBER GREEN Master Mix (PE-Applied Biosystems) and sterile water, for a final volume of 20 μl. The following thermal cycle conditions were used with the Light Cycler 480 (Roche, USA): pre-incubation at 95℃ for 5 min, then 55 cycles of 95℃ for 3 s, 60℃ for 10 s and 72℃ for 30 s, and a final extension at 72℃ for 3 min. Additionally, the expression levels were calculated with the 2^−ΔΔCt^ method for each sample. Data were analyzed using the Office 2010 software, and statistical analyses were conducted with SPSS17.0 software using Duncan’s multiple range test at the *P* < 0.05 level of significance.

## 3. Results

### 3.1 Identification and classification of *SLAC/SLAH* genes in the *Rosaceae, Arabidopsis* and rice

To identify the members of *SLAC/SLAH* gene family in *Rosaceae* and rice, HMM search with the *SLAC/SLAH* gene domain HMM profile (PF03595.14) and BLASTP using *SLAC/SLAH* protein sequences from *Arabidopsis thaliana* as query were performed. We downloaded the *SLAC/SLAH* family of *Arabidopsis*, which contained five *SLAC/SLAH* genes, from the TAIR database. A total of 59 genes was identified in *Rosaceae* and rice using these two strategies. We removed incomplete gene sequences, transcripts of the same gene and redundant sequences. Finally, we identified 30 non-redundant *SLAC/SLAH* genes in the *Rosaceae* and the rice genome (Fig. 1), including four *SLAC/SLAH* proteins identified from pear (*PbrSLAC/SLAH*), five proteins from apple, four proteins from peach, three proteins from strawberry, five proteins from Chinese plum and nine proteins from rice.

**Figure 1.**
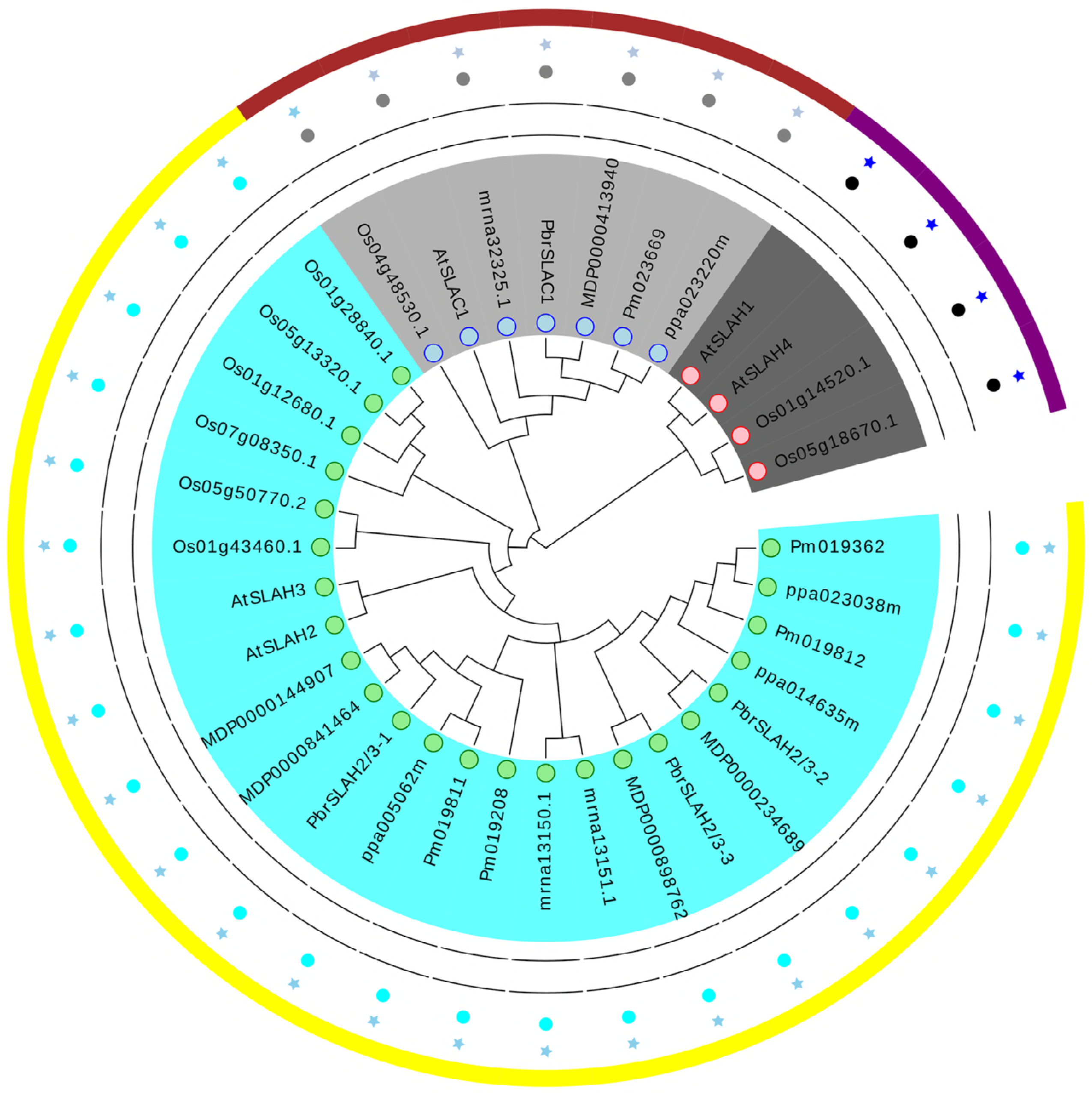
Phylogenetic tree of *Rosaceae*, *Arabidopsis* and rice S-type anion channel genes. The tree was generated using MEGA 6.0 with the Neighbor-Joining method. The proteins clustered into three subgroups. Yellow, brown and purple lines indicate the three subfamilies of the *SLAC/SLAH* proteins.

To classify the *SLAC/SLAH* genes identified in five *Rosaceae* species, *Arabidopsis* and rice, and investigate their evolutionary relationships, we performed phylogenetic analyses using protein sequences encoded by each gene. A phylogenetic tree based on protein sequences, which was constructed using the NJ method, showed that the *SLAC/SLAH* genes from the seven species could be separated into three well-supported clades (bootstrap values of 100%; Fig. 1). A previous study has indicated that the *SLAC/SLAH* gene family was divided into three subfamilies in *Arabidopsis* [9], which is consistent with our grouping. The *PbrSLAC/SLAH* genes are distributed on three chromosomes in pear, with two *PbrSLAC/SLAH* genes detected on chromosome 1 (Table 1). Like the *PbrSLAC/SLAH* genes, the distributions of the *SLAC/SLAH* genes in the other four *Rosaceae* and rice genomes are random.

**Table 1.**
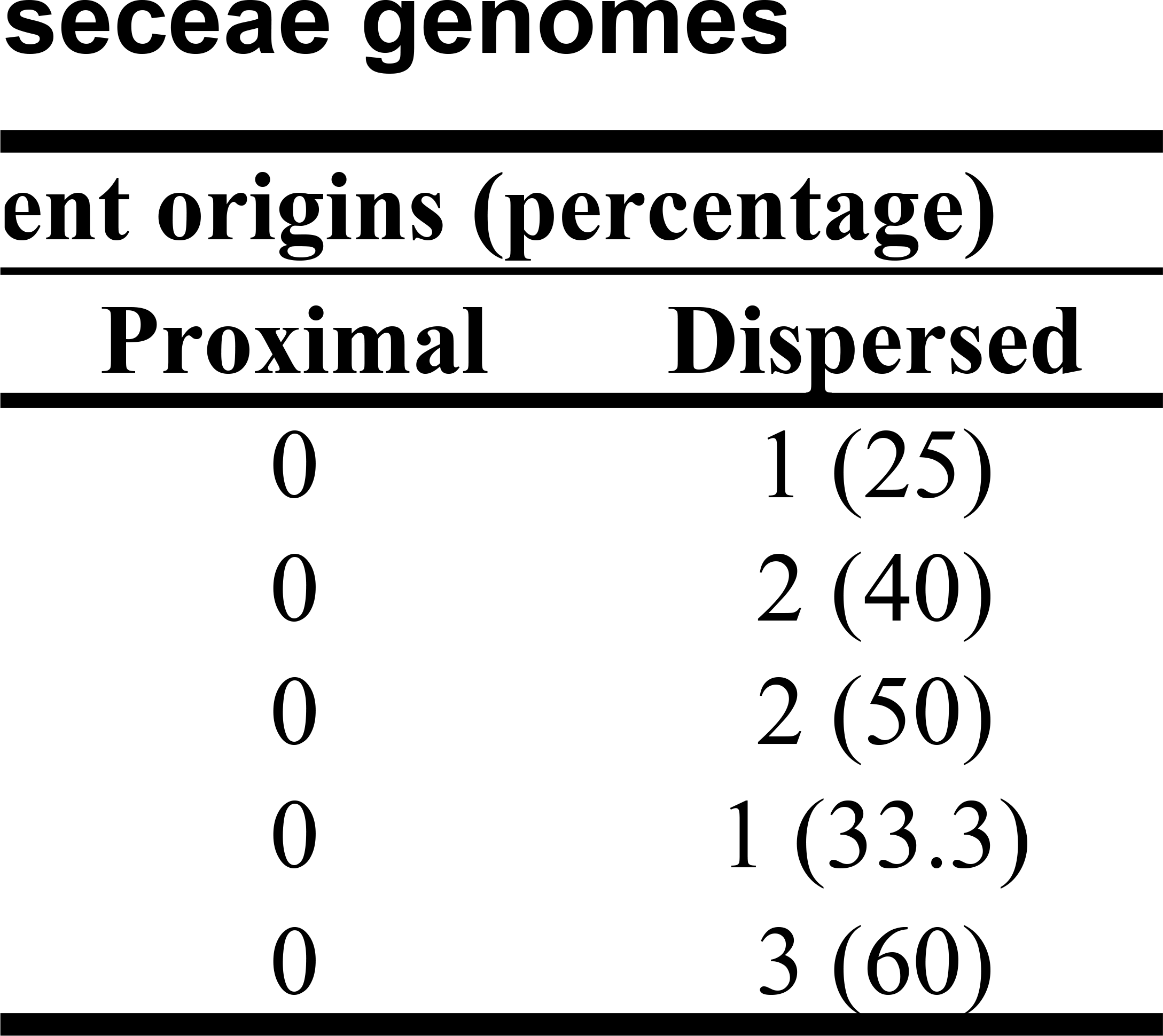
Characteristics of the *SLAC/SLAH* proteins. PI: Isoelectric point RAVY: Grand average of hydropathy Chr: Chromosome

To explore the structural diversity of *SLAC/SLAH* genes in *Rosaceae*, we performed an exon/intron analysis by aligning genomic sequences with their corresponding cDNAs of the *Rosaceae SLAC/SLAH* genes (Fig. 2B). The numbers of exons determined in members of the *SLAC/SLAH* gene family ranged from two to five in *Arabidopsis*, whereas the number of exons ranged from three to five in *Rosaceae,* except for pm019208 from Chinese plum, which only contains two exons (Fig. 2B).

**Figure 2.**
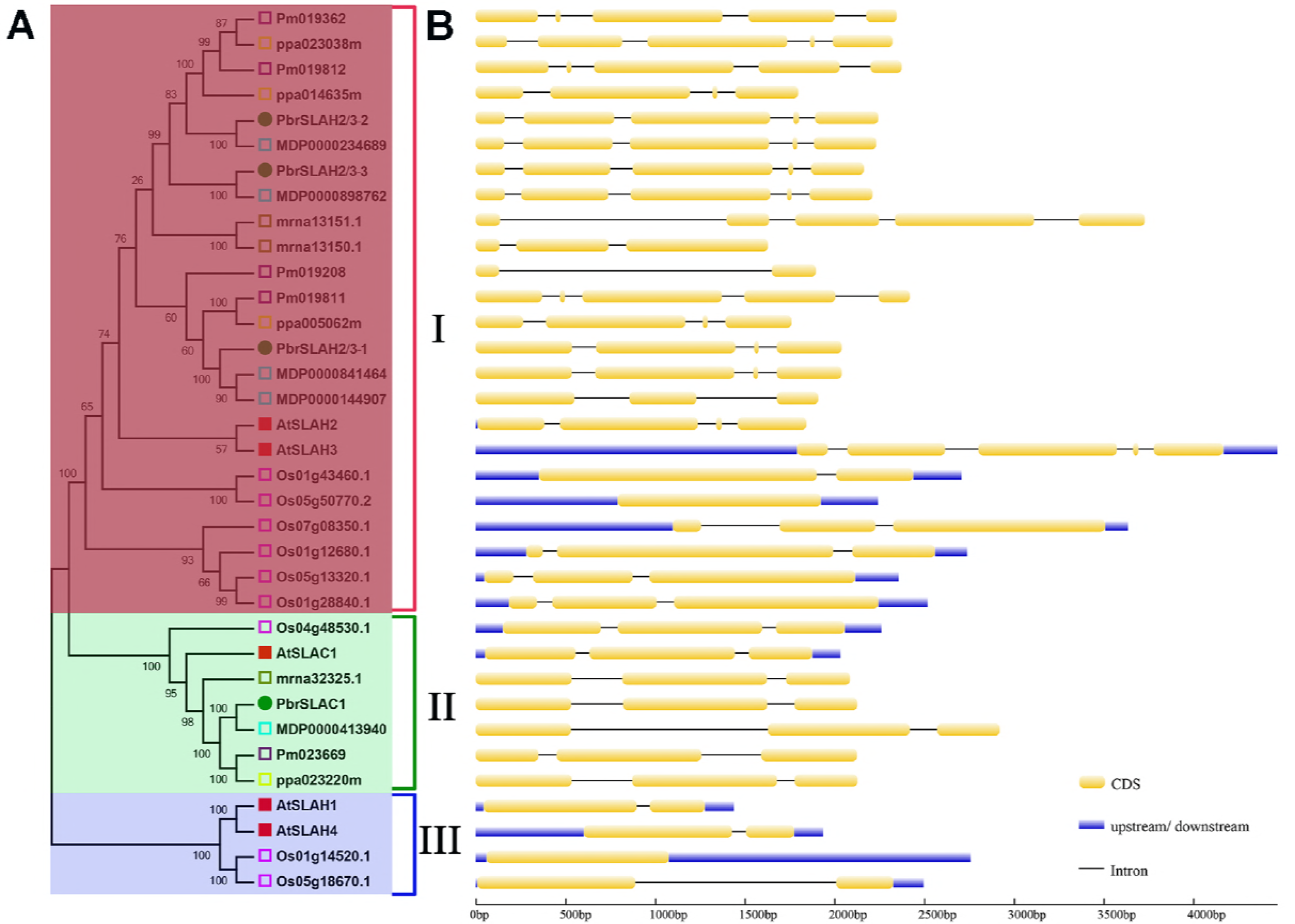
The phylogenetic relationships and schematic diagram for intron/exon structures of *SLAC/SLAH* genes in five *Rosaceae* species, *Arabidopsis* and rice. A. The phylogenetic tree was constructed using the full-length protein sequences of 35 *SLAC/SLAH* genes with the Neighbor-Joining method and 1,000 bootstrap replicates. The numbers beside the branches indicate the bootstrap values that support the adjacent node. B. The yellow boxes, gray lines and blue boxes in the gene structural diagram represent exons, introns and UTRs, respectively. Gene models are drawn to scale as indicated at the bottom.

Most members within the individual groups shared different intron/exon numbers and coding sequence lengths in *Rosaceae*, consistent with the phylogenetic classification of the *SLAC/SLAH* genes (Fig. 2A). Among them, subgroup *SLAH2/3* included members with 4–5 exons in the *SLAC/SLAH* of *Rosaceae*, which is consistent with the numbers of exons (4–5) in the *SLAH2/3* of *Arabidopsis*. Compared with *SLAH2/3*, members of *SLAC1* contained three exons in *Rosaceae*. Interestingly, *SLAH1/4* in group III only have two exons, and no genes were found in the *SLAH1/4* gene subfamily throughout the *Rosaceae* family. This result suggests that different subfamilies have different exon numbers. This may be the results of replication events in the gene family’s evolutionary process, and it also implies that they originated through an evolutionary path separate from that of the other subgroups.

### 3.2 Features of the *SLAC/SLAH* genes in *Rosaceae*, *Arabidopsis* and rice

The characteristics of the 35 *SLAC/SLAH* genes are shown in Table 1. The lengths of the protein sequences varied from 124 to 698 aa, and most of them were from 500 to 698 aa. The protein molecular weights were from 13.48 to 76.24 kD, and the isoelectric points ranged from 6.68 to 10.85 (Table 1). The maximum number of exons in pear *SLAC/SLAH* genes was five, and a similar trend was observed in other *Rosaceae* species (apple, peach, strawberry and plum) and *Arabidopsis* (Table 1). In contrast, the maximum number of exons in rice was three. In pear, the *SLAC/SLAH* genes were located on chromosomes 1, 4 and 13, and two genes (*PbrSLAH2/3-1* and *PbrSLAH2/3-2*) were distributed in chromosome 1. The grand average of hydropathy was positive for all of the proteins in pear, while 82.3% *SLAC/SLAH* protein (two genes) were positive and 17.7% were negative in the other *Rosaceae* species. However, there were three genes with positive values, and two genes with negative values in *Arabidopsis*. In rice, all of the *SLAC/SLAH* proteins had positive values. This indicated that most of the proteins are hydrophobic in the *SLAC/SLAH* gene family, which was similar to the results in *Arabidopsis* (Table 1).

### 3.3 Synteny analysis

Gene duplication modes contributed to the evolution of protein-coding gene families, including WGD, segmental and tandem duplications, and rearrangements at the gene and chromosomal levels [35]. We detected the origins of duplicated genes in the *SLAC/SLAH* gene family in five *Rosaceae* genomes using the MCScanX package. We analyzed the duplication events of *Rosaceae SLAC/SLAH* genes, and each member of the *SLAC/SLAH* gene family was assigned to one of five different types: singleton, WGD or segmental, tandem, proximal, or dispersed. In this study, three types of duplication events drive the expansion of the *SLAC/SLAH* gene family (Table 2). WGD only occurred in pear and accounted for 50%. In addition, only 25% of the *SLAC/SLAH* genes in Chinese white pear and 40% of those in Chinese plum were duplicated and retained from tandem events, compared with 60% in apple, 50% in peach and 66.7% in strawberry. However, the proportions of dispersed *SLAC/SLAH* gene duplications in Chinese plum (60%), peach (50%) and apple (40%) were considerably higher than in pear (25%) and strawberry (33.3%) (Table 2). These results showed that tandem and dispersed gene duplications play critical roles in the expansion of the *SLAC/SLAH* gene family in the *Rosaceae*.

**Table 2.** Numbers of *SLAC/SLAH* genes from different origins in five *Roseceae* genomes. Note: The table shows the total number of *SLAC/SLAH* genes in five *Rosaceae* species, and the number of genes from different duplication events in five *Rosaceae* species.

| Species | No. of *SLAC/SLAH* genes | No. of *SLAC/SLAH* genes from differ |  |  |
| --- | --- | --- | --- | --- |
|  |  | Singleton | WGD | Tandem |
| Chinese white pear | 4 | 0 | 2 (50) | 1 (25) |
| Apple | 5 | 0 | 0 | 3 (60) |
| Peach | 4 | 0 | 0 | 2 (50) |
| Strawberry | 3 | 0 | 0 | 2 (66.7) |
| Chinese plum | 5 | 0 | 0 | 2 (40) |

To explore the evolutionary process behind the *SLAC/SLAH* genes in *Rosaceae*, we investigated the syntenic relationships between pear and six other species, *P. bretschneideri*, *M. domestica*, *P. persica*, *F. vesca*, *P. mume* and *A. thaliana*. The *PbrSLAC/SLAH* genes are distributed on three of the 17 pear chromosomes. Among them, *PbrSLAC1* is on pear chromosome 4, and *PbrSLAH2* and *3* are on pear chromosomes 1 and 13, respectively. *PbrSLAH2/3-1* and *PbrSLAH2/3-2* are co-distributed on chromosome 1. The slow anion channel genes in other *Rosaceae* species and *Arabidopsis* showed random chromosomal distributions (Fig. 3).

**Figure 3.**
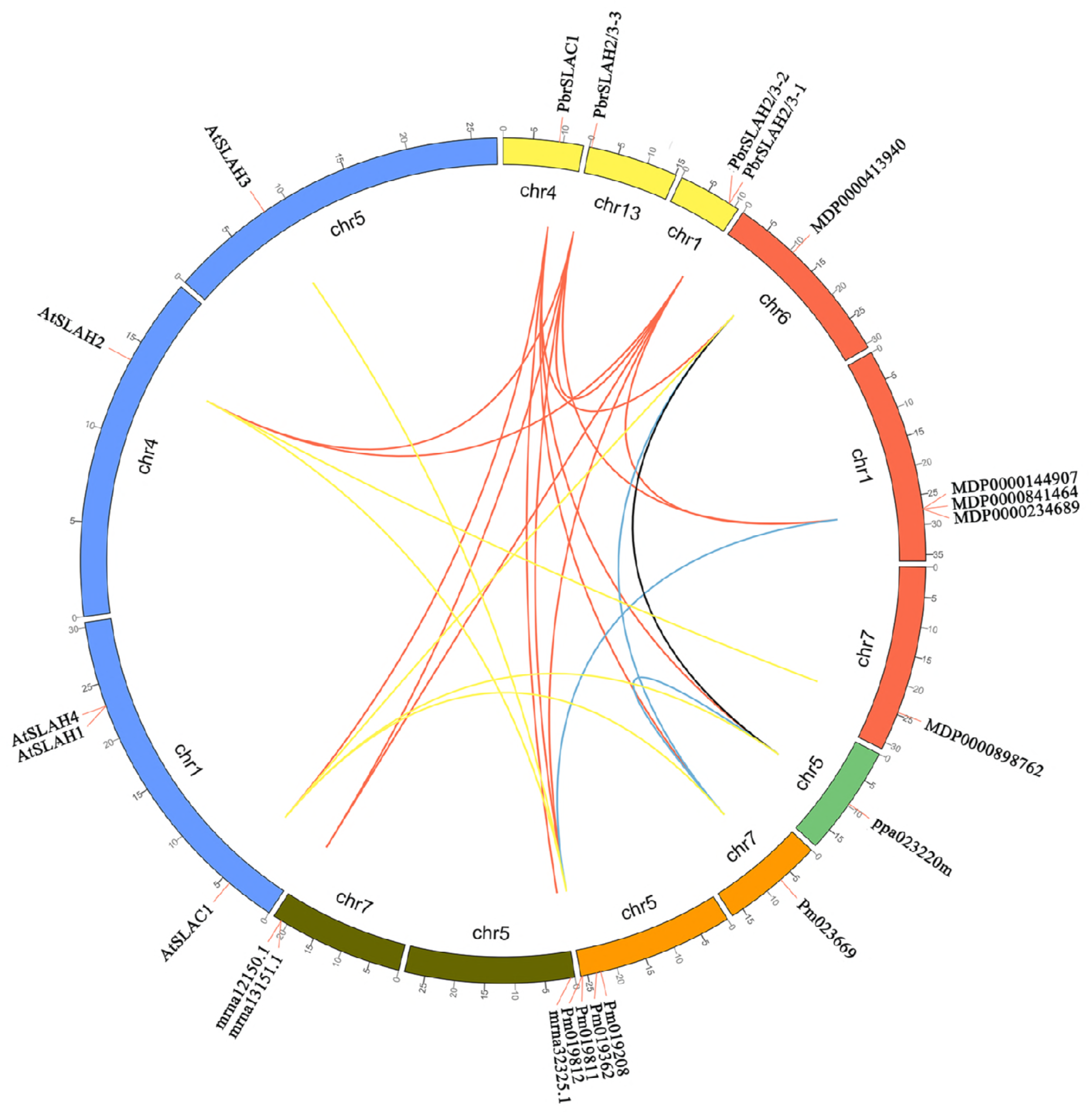
Chromosomal localization of *SLAC/SLAH* genes and syntenic relationships between pear and four *Rosaceae* species between pear and *Arabidopsis*. The circular forms of *Rosaceae* and *Arabidopsis* chromosomes are shown in different colors. The approximate positions of the *AtSLAC/SLAH*, *PbrSLAC/SLAH* and other *SLAC/SLAH* genes are marked with short red lines on the circles. Gene pairs with syntenic relationships are joined by the colored lines.

We detected a syntenic gene pair, *PbrSLAH2/3-1*–*PbrSLAH2/3-3*, which are likely derived from recent WGD events according to the number of synonymous substitutions per synonymous site (Ks) values (Supplementary Table S1, Supplementary Fig. S1). Furthermore, we found one syntenic counterpart in each *Rosaceae* species for three pear *PbrSLAC/SLAH* genes. In *Arabidopsis*, we also found two syntenic orthologs for the three pear *PbrSLAC/SLAH* genes (Supplementary Table S1).

### 3.4 Conserved motif analysis of the *SLAC/SLAH* gene family in *Rosaceae* species

To further identify and examine the conserved motifs of the *SLAC/SLAH* gene family in *Rosaceae*, and the MEME program was used in this study. As shown in Fig. 4A and B, 15 conserved motifs with low E values were found (Supplementary Table S2). The number of motifs present in the *SLAC/SLAH* protein sequences was variable. Among them, 15 motifs shared in most of *Rosaceae SLAC/SLAH* gene families, except strawberry, which may result from strawberry having lost a number of *SLAC/SLAH* functional motifs during the evolutionary process of the *SLAC/SLAH* genes (Fig. 4A and B). In pear, motifs 1, 2, 3, 4, 5, 6, 7, 8, 9 and 13 are common functional motifs in genes that form the low anion channel of pear. Motifs 10, 11, 12, 14 and 15 are distributed in different slow anion channel genes of pear, which may determine its specific functions in different organizations. In apple, motifs 1, 3, 4 and 8 were present in all of the *MdSLAC/SLAH* gene families. Motifs 1, 2, 3, 4, 5, 6, 7, 8 and 9 are commonly distributed in the *PpSLAC/SLAH* gene family of peach. In strawberry, motif 15 was not detected in the *FvSLAC/SLAH* gene family. Except for the pm019208 gene, which contains only motifs 4 and 7, the other genes contain most of the *SLAC/SLAH* gene family motifs of plum. Our results indicated that these differences represent the evolutionary relationship of the *SLAC/SLAH* gene family in *Rosaceae* species, suggesting that the presence of motifs 1, 3, 4, 8 and 10 may form the conserved functional motif of the *SLAC/SLAH* gene family of the *Rosaceae*.

**Figure 4.**
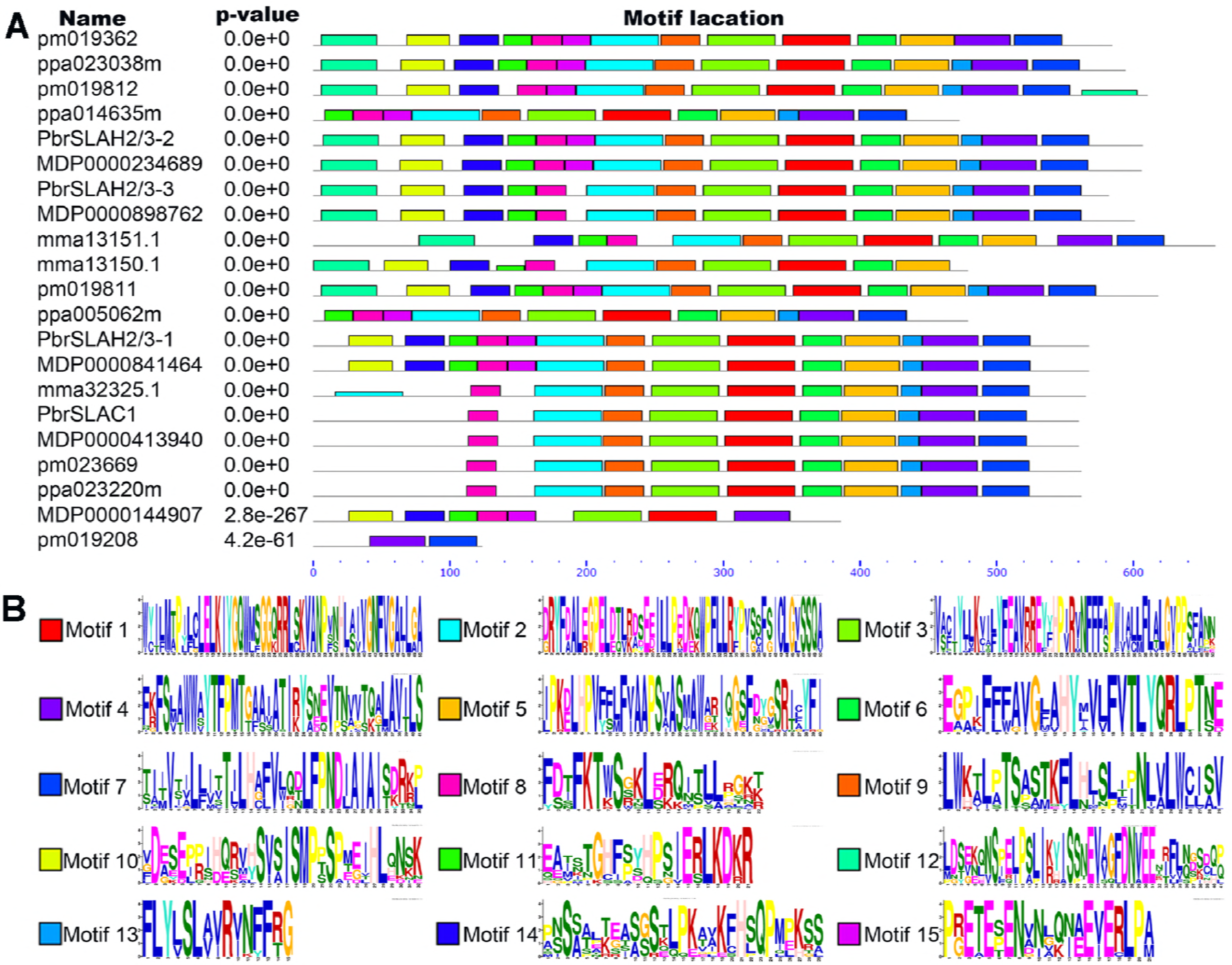
Motifs and sequence logos identified by MEME tools in the *SLAC/SLAH* genes of five *Rosaceae* species. A. In total, 15 motifs (motifs 1 to 15), shown in different colors, were identified. Motifs locations and combined p-values are shown in the figure. The compositions of motifs in the *SLAC/SLAH* genes of the five *Rosaceae* species were identified using the MEME tool. The lengths and positions of the colored blocks correspond to the lengths and positions of motifs in the individual protein sequences. The scale indicates the lengths of the proteins as well as the motifs. B. Over-represented motifs in the five *Rosaceae* species were identified using the MEME tool. The stack’s height indicates the level of sequence conservation. The heights of the residues within the stack indicate the relative frequencies of each residue at that position.

### 3.5 Subcellular localization of the *SLAC/SLAH* genes in pear and *Arabidopsis*

In *Arabidopsis*, the *SLAC/SLAH* genes were those of the S-type anion channel located at the plasma membrane of guard cells. The subcellular localization of only one gene was analyzed in a previous report in *Arabidopsis*. Therefore, to further verify that the *SLAC/SLAH* genes of *Rosaceae* and *Arabidopsis* were all localized on the plasma membrane, we performed the structure analyses of *PbrSLAC/SLAH* genes of pear using ExPASy software. The sequence analysis showed that the *SLAC/SLAH* proteins contain 10 trans-membrane regions (Fig. 5A–D), indicating that they may be localized on the membrane. To confirm on which membrane they were located, four *PbrSLAC/SLAH* genes were cloned from roots and leaves of pear, and five *AtSLAC/SLAH* genes were cloned from roots and leaves of *Arabidopsis*. The open-reading frame of *PbrSLAC/SLAH* was fused to the N-terminus with the GFP protein under the control of the CaMV 35S promoter. The fusion proteins (*PbrSLAC/SLAH*-GFP) and control (35S-GFP alone) were separately transformed into protoplast cells of *Arabidopsis*. Microscopic observations showed that the green fluorescence was distributed in the whole cells when the control plasmid was used, whereas green fluorescence was only detected on the plasma membrane when the vectors contained *PbrSLAC/SLAH*-GFP (*PbrSLAC1*-GFP, *PbrSLAH2/3-1*-GFP, *PbrSLAH2/3-2*-GFP and *PbrSLAH2/3-3*-GFP) (Fig. 5E–I). The plasma membrane localization of *AtSLAC/SLAH* genes was also confirmed in *Arabidopsis* mesophyll protoplasts (Supplementary Fig. S2). Microscopic visualization showed that the control 35S-GFP was distributed throughout the whole cell (Supplementary Fig. S2A), whereas the *AtSLAC/SLAH*-GFP (*AtSLAC1*-GFP, *AtSLAH1*-GFP, *AtSLAH2*-GFP, *AtSLAH3*-GFP and *AtSLAH4*-GFP) fusion protein was observed exclusively on the plasma membrane (Supplementary Fig. S2B–F). Thus, the *PbrSLAC/SLAH* genes may have a similar subcellular location pattern as the *Arabidopsis AtSLAC/SLAH* genes, which were all plasma membrane protein.

**Figure 5.**
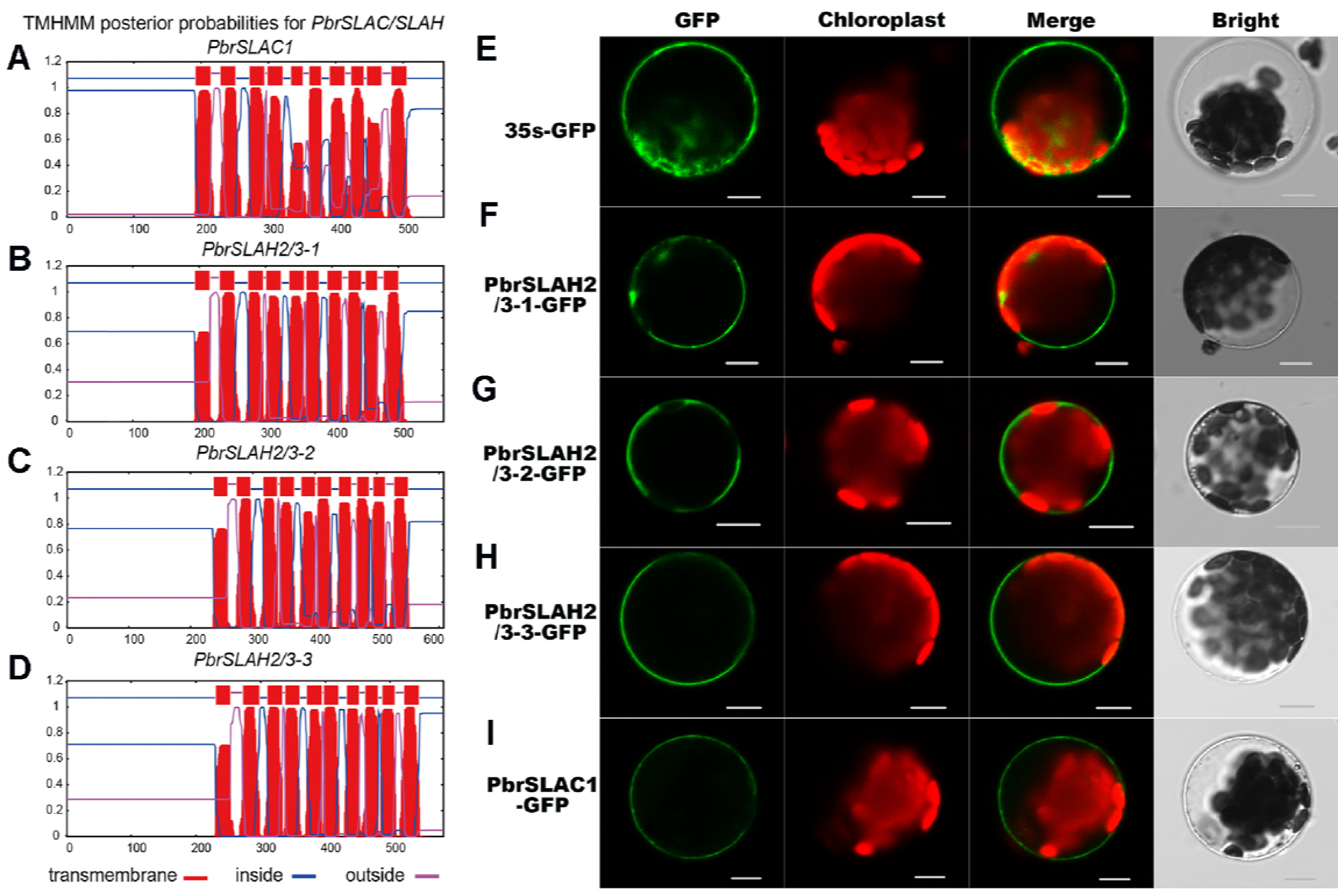
Multi-transmembrane pattern and subcellular localization of *PbrSLAC/SLAH* in pear. A–D. Multi-transmembrane pattern of *PbrSLAC/SLAH* in pear. E–I. The selected *PbrSLAC/SLAH* genes were cloned from pear and used to construct CaMV35S::*SLAC/SLAH*-GFP vectors in which GFP was fused at the C termini. The four *PbrSLAC/SLAH*-GFP fusion proteins (*PbrSLAC1*-GFP, *PbrSLAH2/3-1*-GFP, *PbrSLAH2/3-2*-GFP and *PbrSLAH2/3-3*-GFP) as well as 35S-GFP as the control were independently transiently expressed in *Arabidopsis* Col-0 protoplasts and imaged under a confocal microscope. The merged images include the green fluorescence channel (first panels) and the chloroplast autofluorescence channel (second panels). The corresponding bright field images are shown on the right. Bar = 10 μm

### 3.6 Expression levels of the *SLAC/SLAH* genes in pear

To analyze the expression patterns of the *SLAC/SLAH* genes of different tissues in *Rosaceae*, several candidate genes were selected for quantitative RT-PCR (qRT-PCR) (Fig. 6A). The transcript accumulation patterns were analyzed in roots, stems, leaves, flowers, fruit, pollen grains and pollen tubes of pear cultured for 5 h. qRT-PCR results indicated that the *PbrSLAC/SLAH* gene expression pattern corresponded with the slow anion channel-related genes in *Arabidopsis*. Most of the *PbrSLAC/SLAH* genes in pear were mainly expressed in root and leaf. For example, *PbrSLAC1* and *PbrSLAH2/3-1* showed preferential expression levels in leaf and root. However, the expression level of *PbrSLAC1* in flower was the third largest of all plant tissues, after root, which had very low expression levels in other tissues. *PbrSLAH2/3-1* was mainly expressed in root, in addition to its expression in leaf, fruit and pollen grain. However, *PbrSLAH2/3-2* and *PbrSLAH2/3-3* showed predominant transcript accumulations in root, while *PbrSLAH2/3-2* was lowly expressed in stem and leaf. *PbrSLAH2/3-3* also showed low expression levels in stem, leaf, flower, fruit and pollen.

**Figure 6.**
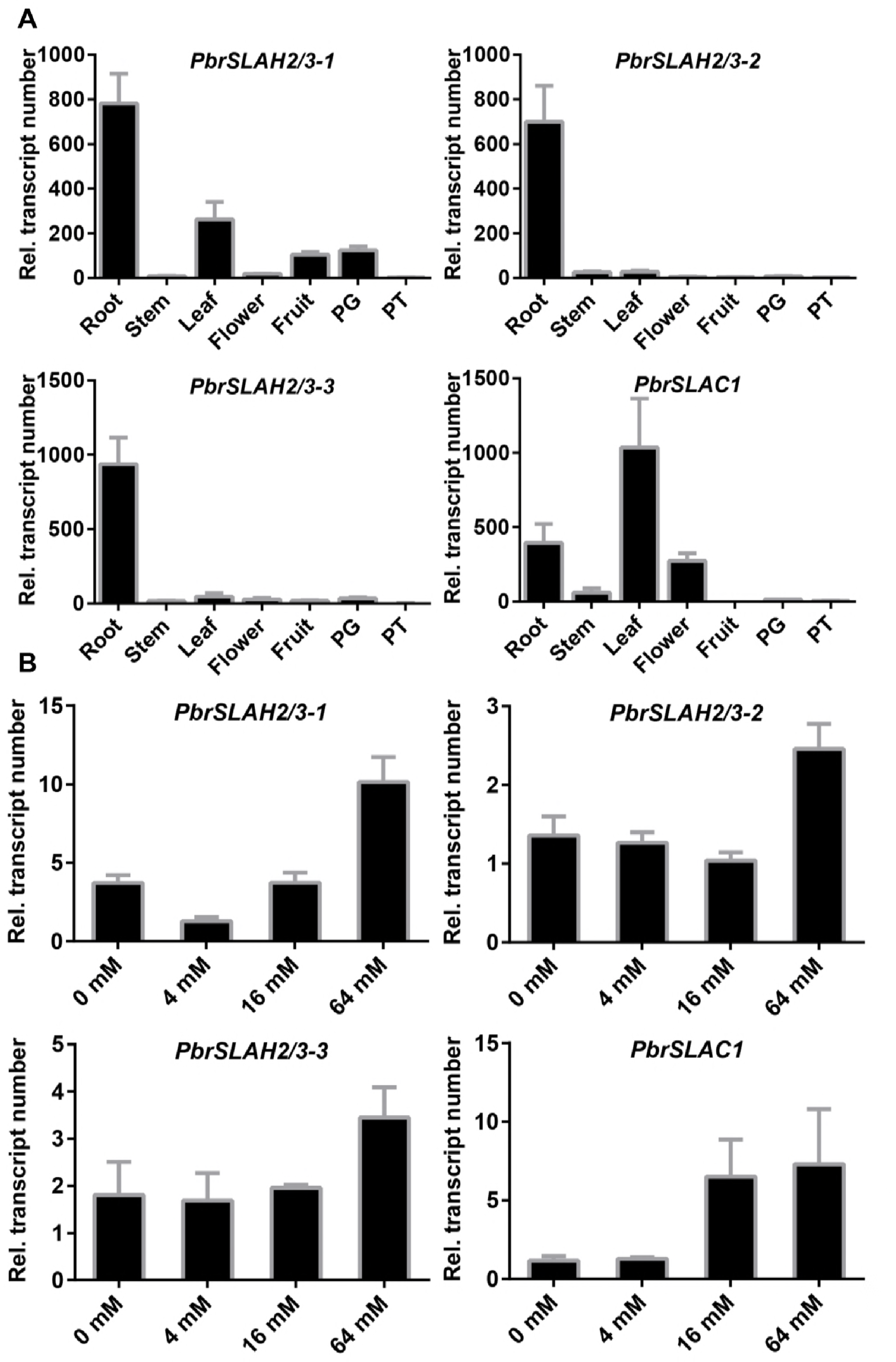
Expression analyses of the four *PbrSLAC/SLAH* genes using qRT-PCR. A: Expression analyses of the four *PbrSLAC/SLAH* genes using qRT-PCR in different tissues of pear. Total RNA was extracted from roots, stems, leaves, flowers, fruit, pollen grains and pollen tubes cultured for 5 h. For each gene, the relative expression levels were obtained by normalization with pear UBQ. B: Expression analyses of the four *PbrSLAC/SLAH* genes in plant root as assessed by qRT-PCR after nitrate ion treatments. Total RNA was extracted from pear root. For each gene, the relative expression levels were obtained by normalization with pear Actin. The error bars indicate standard deviations. The data are shown as mean values ± SDs.

### 3.7 Nitrate transport for the *SLAC/SLAH* gene family in pear

To investigate the mechanisms of slow anion channel uptake and the transport of nitrate ions in the *Rosaceae* species, we selected the pear as the representative. The expression of slow anion channels in roots of pear seedlings under different concentrations of nitrate ions was determined. In this study, the expression level of the *PbrSLAC1* gene increased with the elevation of nitrate ion concentrations in the root of pear, as shown in Fig. 6B. However, the expression level of *PbrSLAH2/3-1* decreased when the concentration of nitrate ions was 4 mM, and then increased as the nitrate ion concentration increased. In addition, the expression levels of *PbrSLAH2/3-2* and *PbrSLAH2/3-3* abruptly increased when the nitrate concentration was excessively (64 mM) increased. Thus, the slow anion channel may function in absorbing and transporting nitrate ions in pear. However, most of the slow anion channels play roles in the excess nitrate supply. This may be analogous to the function of *AtSLAH2* in *Arabidopsis*, which is mainly involved in transporting excess nitrate ions from root to shoot (Fig. 6B).

## 4. Discussion

The identification and functions of homologous *SLAC/SLAH* genes has been well studied in some model plants, such as *Arabidopsis* [36], rice [10] and poplar [11]. However, research on the *SLAC/SLAH* genes family is limited in the *Rosaceae* species. Although the pear genome size (512 Mbp) is larger than that of *Arabidopsis* (130 Mbp), rice (430 Mbp) or poplar (483 Mbp) [37], only four *SLAC/SLAH* genes were identified in pear (*PbrSLAC/SLAH*) compared with five *SLAC/SLAH* genes in *Arabidopsis*. The rare homologous genes of *SLAC/SLAH* found in pear and other plants suggested that slow anion channel genes were resistant to duplication though the rampant WGD events that occurred in plant genomes. These *SLAC/SLAH* genes, including *SLAC1*, *SLAH2/3* and *SLAH1/4*, were classified into three subfamilies, I, II and III. This is consistent with the classification of *Arabidopsis SLAC/SLAH* genes [9]. However, subfamily *SLAC1* members only contain five genes in the *Rosaceae*, and no member has been found in subfamily III. Most of the members are clustered in subfamily I. We found some evidence for the absence of subfamily III members in the *Rosaceae* by investigating the numbers of introns and exons. Three pivotal mechanisms contribute to the inactivation of genes: exon/intron gain or loss, exonization/pseudo-exonization and insertion/deletion [38].

Pear and apple are thought to have undergone a recent lineage-specific WGD, while peach, strawberry and Chinese plum did not experience this event [39]. We found that most group I members contain 4–5 exons, while only a small set 1–2 exons, which may be the result of fragment loss during gene duplication. Members of group II contain three exons, and group III members contain 1–2 exons. Thus, the *SLAC/SLAH* family of genes has evolved various intron–exon organizations over its long evolutionary history. The intron–exon divergence among *SLAC/SLAH* family genes might result in functional diversity. Group I contained the most exons, and this may indicate that more intron insertion events have occurred in the exons of group I genes, which contributed to their divergent functional roles in the different organs and tissues. Therefore, in the *Rosaceae* species, slow anion channel gene function is mainly concentrated in group I. In addition, we also investigated the conserved motifs of *SLAC/SLAH* genes and determined the putative protein localization as well as their collinearity relationships. The conserved motif analysis suggested that the presence of motifs 1, 3, 4, 8 and 10 may indicate a conserved functional motif in the *SLAC/SLAH* gene family of *Rosaceae*. The *SLAC/SLAH* gene family plays important roles in adjusting pollen tube growth, stomatal closure and the transport of root anions [4, 40, 41]. *SLAC/SLAH* genes have a wide range of functions in plant growth and development. For example, *AtSLAC1* encodes a plasma membrane-localized protein that is highly permeable to malate and chloride in *Arabidopsis* [42], and our experimental results also support this observation (Supplementary Fig. S2). The real-time fluorescence quantitative PCR showed that *PbrSLAC1* is predominantly expressed in leaves. Its homologous gene was activated by the stress hormone ABA, ozone, CO_2_, calcium, light/dark transitions and reduced humidity levels, and is involved in the early steps leading to stomatal closure [15, 43]. Furthermore, *OsSLAC1* can be phosphorylated and activated by *OsSAPK8* in rice, which is a nitrate-selective anion channel [10]. *AtSLAC1* in *Arabidopsis* was co-distributed in group II, which contains the *Rosaceae SLAC1* genes. These genes are very similar to each other in structure, subcellular localization and expression pattern. Therefore, we speculated that the *SLAC1* genes from *Rosaceae* may function in the leaves, and be involved in responding to changes in the external environment and abiotic stresses that control the stomatal switch. Most members of *Rosaceae* species are found in this group I, including *SLAH2* and *SLAH3*. Therefore, members of this subfamily may play important roles in the *Rosaceae*. *AtSLAH3*, which is closely related to *AtSLAC1*, appears to have an overlapping function with *SLAC1* in guard cells [42]. Compared with *AtSLAC1*, *AtSLAH3* exhibits a stronger selectivity for nitrate over chloride (20-fold); therefore, it can be considered a nitrate efflux channel [16, 44]. *AtSLAH3* is also located on the plasma membrane (Supplementary Fig. S2), as previously reported [36]. This indicates that *SLAH3* plays an important role in the transport of membrane anions. However, in the *Rosaceae*, the *SLAH2* and *SLAH3* genes showed high expression levels in root. Therefore, we explored the functional roles of SLAH2 and SLAH3 proteins in the nitrate transport of root. *AtSLAH3* functions as an outwardly rectifying anion channel [16], alleviates NH_4_^+^ toxicity that is dependent on the nitrate [9] and adjusts pollen tube growth through the calcium-dependent protein kinases in *Arabidopsis* [21]. In addition, the functions of *SLAH3* have been well studied in many other species. For example, *PttSLAH3* may form a secretory cell with a channel that controls long-term nectar secretion in poplar [11]. *SLAH2* is the closest homolog of *SLAH3* and is also expressed in root tissues. Furthermore, *AtSLAH2* is mainly expressed in the stele of the root, which are the cells that immediately surrounding the vasculature. These cells determine the anion composition of the sap flow between roots and shoots [45], and the protein is mainly permeable to NO_3_^−^ (with a NO_3_^−^/Cl^−^ permeability ratio of 82) [46]. Thus, *SLAH2* may have evolved to facilitate root–shoot nitrate transport in the *Rosaceae*. *SLAH1* and *SLAH4* belong to group III; however, no members of this subfamily were found in the *Rosaceae*. Furthermore, the gene function of *AtSLAH1* is poorly understood, but it is mainly expressed in root. *AtSLAH1*, through the formation of *SLAH1/SLAH3* heteromers, facilitates Cl^−^ efflux by rendering *AtSLAH3* nitrate phosphorylation independent in *Arabidopsis* [47]. In addition, the role of *AtSLAH4* is similar to that of *AtSLAH1*, and they play important roles for chloride ion and nitrate transport in root. These results lay a foundation to further study the functions and regulation of genes related to the S-type anion channel by analyzing the structures and characteristics of their expression levels using bioinformatics methods in the *Rosaceae*.

## Conclusion

In short, few bioinformatics analyses of the *SLAC/SLAH* family have been reported for the *Rosaceae*; therefore, the functions and characteristics of most *SLAC/SLAH* genes remain unclear. However, evidence indicates that *SLAC/SLAH*s play important roles in responses to stress signaling, and growth and development. In this study, 21 full-length *SLAC/SLAH* genes were identified in *Rosaceae* genomes, with the pear genome containing 4 *SLAC/SLAH* genes. The structural characteristics of the encoded proteins, phylogenetic analysis, expression analyses and comparisons with homologues from *Arabidopsis* and rice provide a framework for further analyses of the *Rosaceae SLAC/SLAH* genes to define their biological functions and pathways during stress responses, as well as during growth and development. An expression analysis showed that pear *SLAC/SLAH* genes respond to different concentrations of nitrate ions and were expressed in different tissues. Moreover, the *PbrSLAC/SLAH* subcellular localizations provided information regarding their functions. Comparisons of these genes in *Rosaceae* and *Arabidopsis* genomes, and their protein characteristics, provided insights into the functions of these less well-studied genes by better understanding their homologs. Based on previous experimental data and the current bioinformatics analysis, we predicted that the *SLAC/SLAH* gene family might participate in responding to nitrate and chloride transport, in stomatal closure regulation, in pollen tube growth regulation, and in mediating the cross-talk between signaling pathways. Our results form a foundation for further studies that will examine the structures and functions of the *SLAC/SLAH* gene family in the *Rosaceae*.

## Acknowledgements

This work was supported by the National Key Technology R & D Program of the Ministry of Science and Technology of China (2014BAD16B03-4), the Fundamental Research Funds for the Central Universities (KYTZ201602) and the China Postdoctoral Science Foundation (2017M621760).

## Supplementary materials

**Supplementary table S1.**
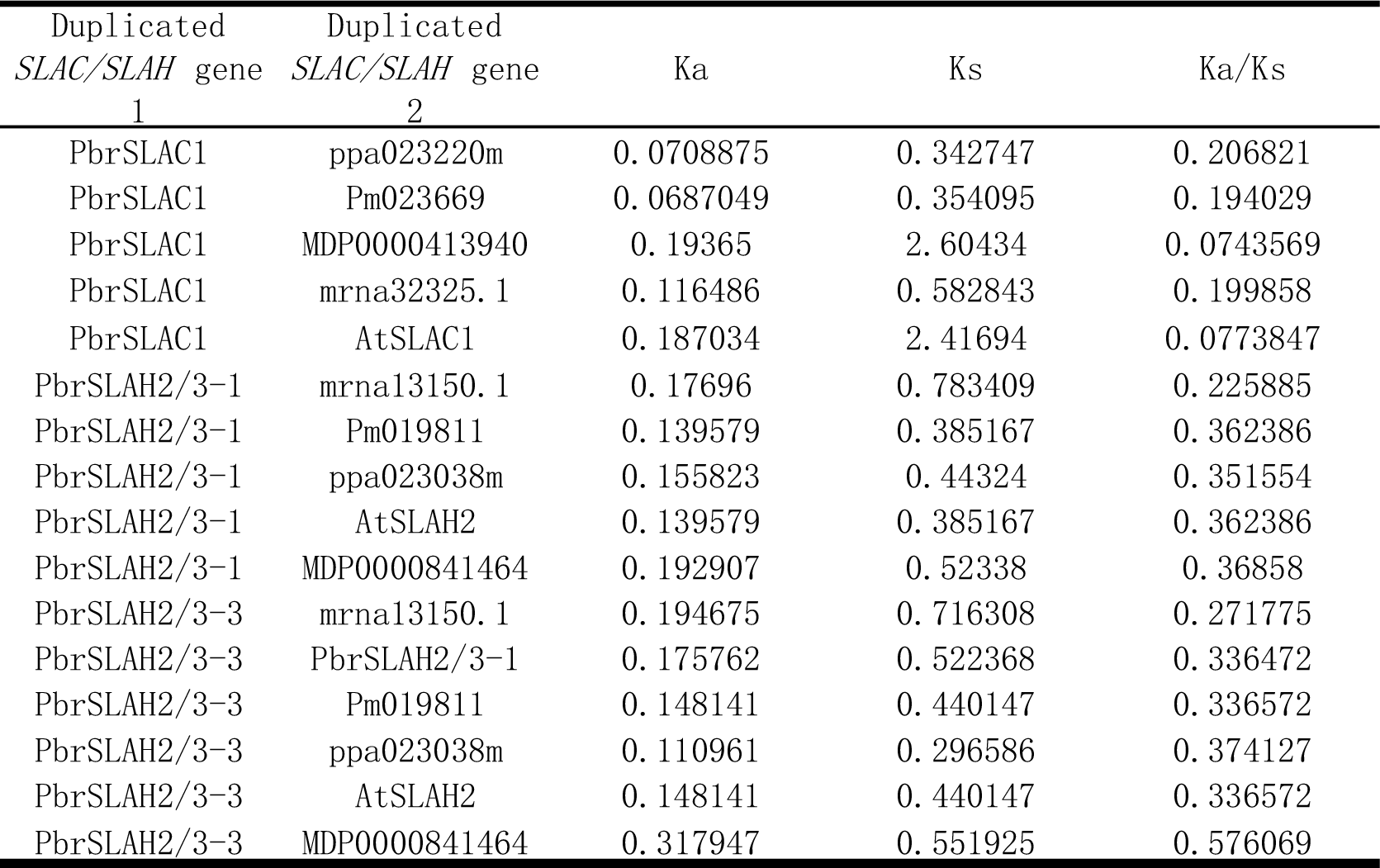
Synteny analysis of *SLAC/SLAH* gene regions in *Rosaceae* species and the *Arabidopsis* genome.

**Supplementary table S2. Motif distribution in *SLAC/SLAH* families.**

**Supplementary table S3. Primers used in this study.**

**Supplementary figure S1.**
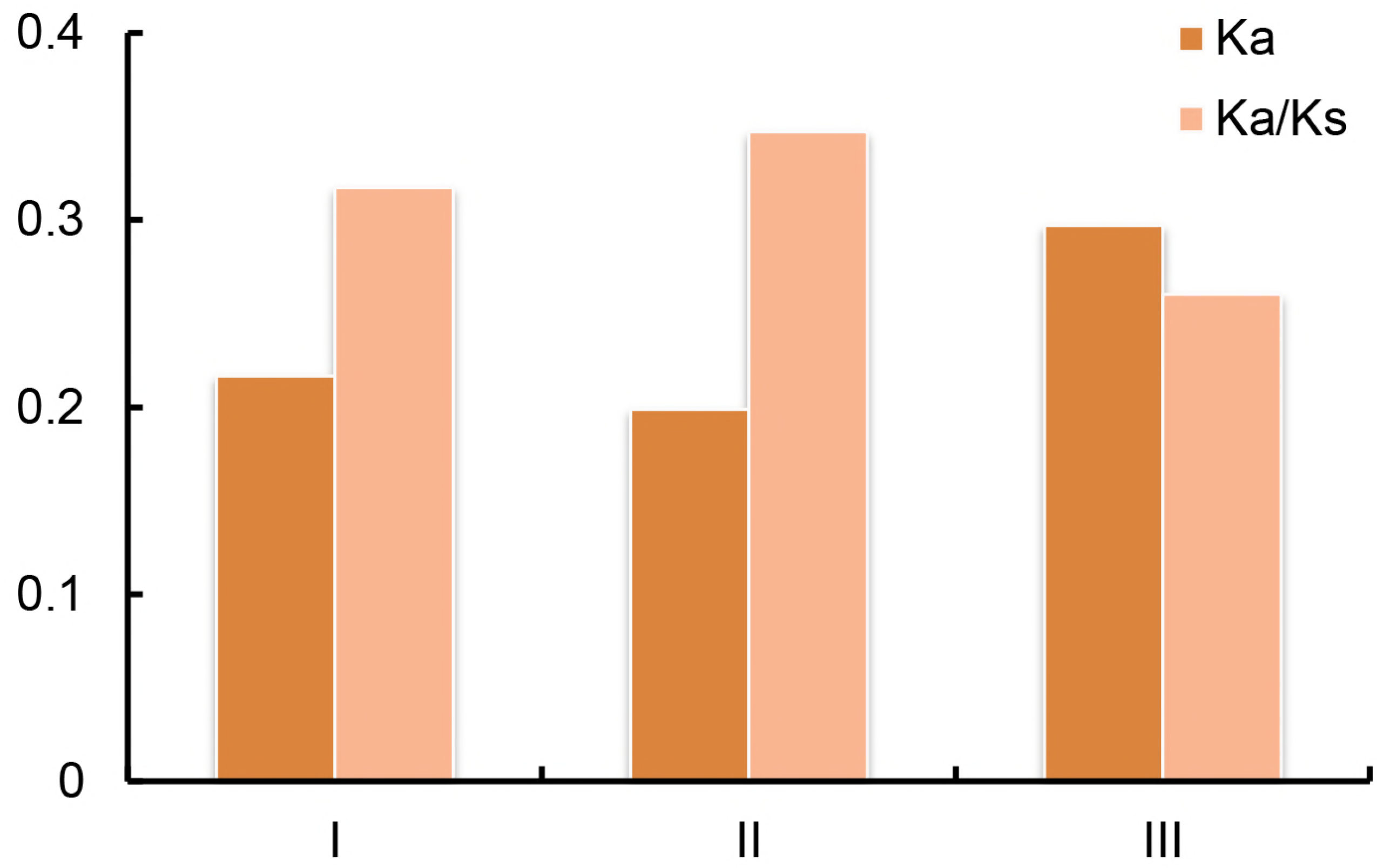
Distribution of mean Ka and Ka/Ks values of *SLAC/SLAH* genes in *Rosaceae* species.

**Supplementary figure S2.**
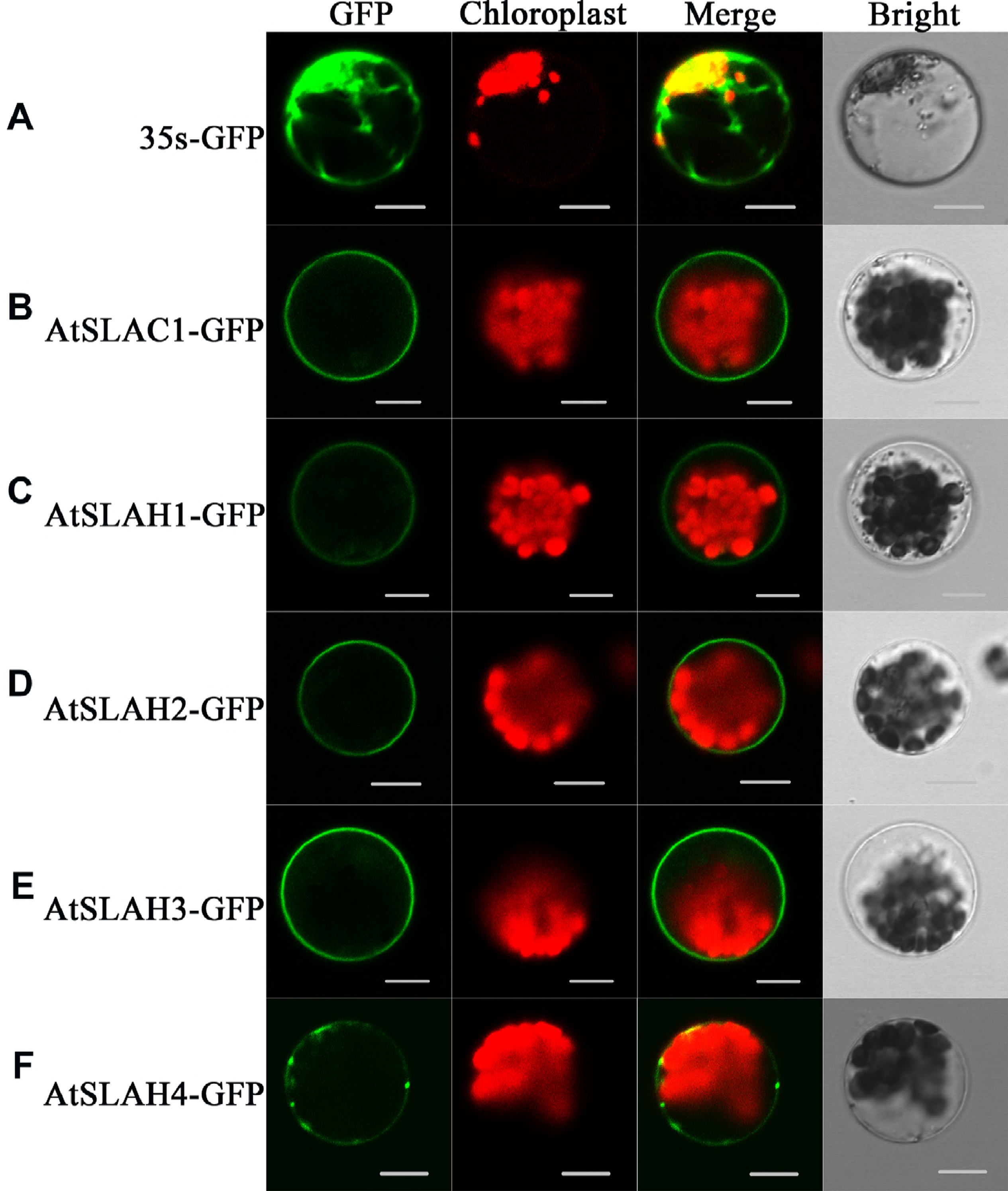
Subcellular localization of five *AtSLAC/SLAH* genes.

